# Mechanical Loading Induces the Radial Growth of Myofibrils and Myofibrillogenesis via an mTORC1-Dependent Mechanism

**DOI:** 10.64898/2026.05.18.725456

**Authors:** Corey GK. Flynn, Ramy K. A. Sayed, Anthony N. Lange, Wenyuan G. Zhu, Troy A. Hornberger

**Affiliations:** Department of Comparative Biosciences, University of Wisconsin-Madison, Madison, WI, USA; School of Veterinary Medicine, University of Wisconsin-Madison, Madison, WI, USA; Department of Anatomy and Embryology, Faculty of Veterinary Medicine, Sohag University, Sohag, Egypt

**Keywords:** mTORC1, Raptor, Skeletal muscle, Myofibril, Myofibrillogenesis, Ultrastructure, Hypertrophy, Mechanical Overload, Sarcomere

## Abstract

Increased mechanical loading induces skeletal muscle growth and, at the ultrastructural level, promotes myofibrillogenesis and the radial growth of myofibrils. However, the mechanisms regulating these ultrastructural adaptations are not known. Here, we sought to determine whether the mechanistic target of rapamycin complex 1 (mTORC1) regulates these processes. To accomplish this, muscle-specific, tamoxifen-inducible raptor knockout (iRAmKO) mice were used to inhibit signaling through mTORC1, and growth was induced with a model of chronic mechanical overload (MOV). Using a next-generation fluorescence imaging pipeline for ultrastructural analyses, we found that mTORC1 is a critical regulator of the myofibrillogenesis and radial growth of myofibrils that occur in response to MOV. Together with other recent advances in the field, we propose a model in which mTORC1 acts as a gatekeeper that permits the retention, rather than the synthesis, of proteins that drive the ultrastructural adaptations.

## Introduction

Skeletal muscle comprises approximately 45% of total body mass and plays major roles in numerous bodily functions, such as voluntary movement, respiration, whole-body metabolism, and regulation of body temperature ^1-3^. As adults age, they will lose 30-40% of their muscle mass by the age of 80 ^1,3^. Importantly, this loss of muscle mass is correlated with increased prevalence of chronic disease, decreased independence, and diminished quality of life ^4-6^. Accordingly, defining the mechanisms that regulate the maintenance and growth of skeletal muscle is a critical objective for improving healthspan.

Many studies have demonstrated mechanical loading to be a potent regulator of skeletal muscle mass. Reductions in loading, such as those caused by bed rest or immobilization, result in decreased muscle mass (i.e., atrophy) while increased loading, such as that which occurs during resistance exercise, promotes an increase in muscle mass (i.e., hypertrophy) ^7-12^. This hypertrophic response is at least in part due to an increase in the cross-sectional area (CSA) of the muscle fibers (i.e., radial growth of muscle fibers) ^13,14^. However, relatively little is known about the structural adaptations that drive this response. For instance, only a few studies have attempted to define the adaptations that occur at the level of the myofibrils, which contain the contractile elements of muscle fibers and occupy ∼80% of the fiber volume ^13,15,16^. This gap in knowledge largely reflects the technical hurdles that are associated with the imaging of myofibrils. Specifically, electron microscopy has been the gold standard method for the imaging of myofibrils, but it is highly time- and cost-intensive ^17^. Further, electron microscopy generates relatively low-contrast images that are not conducive to automated myofibril tracing. As a result, the use of electron microscopy requires manual tracing, and the tracing of enough myofibrils for even a modestly powered study could take hundreds to thousands of hours of labor, essentially preventing researchers from pursuing such studies ^18,19^.

To overcome these limitations, we recently developed a technique called fluorescence imaging of myofibrils with image deconvolution (FIM-ID), a light-microscopy-based approach that integrates standard immuno-labeling with automated myofibril quantification ^18^. Using this method, we demonstrated that myofibril measurements obtained with FIM-ID are indistinguishable from manual measurements derived from electron microscopy. Moreover, the use of FIM-ID enabled us to show that chronic mechanical overload (MOV) induces robust myofibrillogenesis (i.e., an increase in the number of myofibrils per fiber) as well as radial growth of individual myofibrils (i.e., an increase in myofibril CSA) ^18^. However, the mechanisms that regulate these distinct forms of growth remained unclear. Nonetheless, previous studies have shown that signaling through the mechanistic target of rapamycin complex 1 (mTORC1) is necessary for the radial growth of muscle fibers that occurs in response to MOV ^7,20-22^. Based on this point, we reasoned that mTORC1 plays an equally critical role in the induction of myofibrillogenesis and the radial growth of myofibrils. Hence, this study was designed to directly interrogate the role of mTORC1 in these fundamental processes.

## Results

### Mechanical Overload Induces Radial Growth of Muscle Fibers via an mTORC1-Dependent Mechanism

To identify the growth-related adaptations that are mediated through an mTORC1-dependent mechanism, we utilized our previously described skeletal muscle-specific, tamoxifen-inducible raptor knockout mice (iRAmKO) and control littermates ^7^. Specifically, at 6 weeks of age, all mice were treated with tamoxifen for 5 consecutive days. Two weeks after completing the treatment, the mice underwent either an MOV or sham surgery (Fig. 1A). Notably, the MOV surgery involved the removal of the distal third of the gastrocnemius muscle, while leaving the plantaris (PLT) and soleus muscles intact. This is noteworthy because, as previously demonstrated, this approach does not elicit the signs of damage/regeneration that are commonly observed with the traditional synergist ablation model of MOV, in which both the gastrocnemius and soleus muscles are removed ^7,23^. As such, the use of MOV allowed for the assessment of mechanical overload-induced adaptations while minimizing the confounding influences of damage/regeneration. It also bears mentioning that both male and female mice were included in this study, and none of the primary outcomes showed a significant interaction between the effects of sex, genotype, and MOV (Fig. S1). However, the baseline values for many of the variables (e.g., whole muscle CSA, fiber CSA, and myofibrils/fiber) were significantly lower in the muscles from female mice. Thus, to correct this, the values for each animal were expressed relative to the mean of their sex-matched controls.

**Figure 1.**
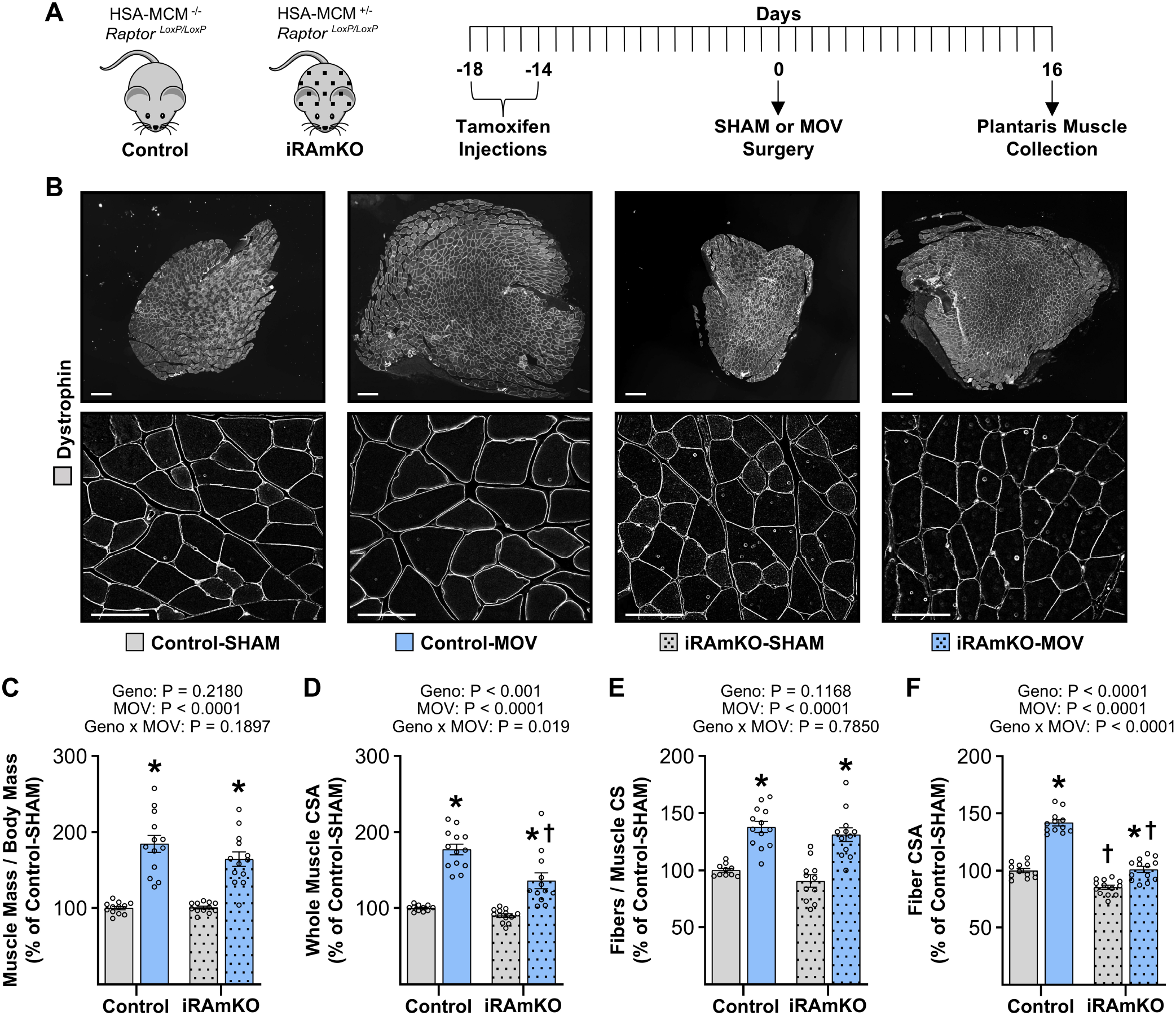
Mechanical Overload Induces Radial Growth of Muscle Fibers via an mTORC1-Dependent Mechanism. (A) Schematic of the experimental design. Briefly, male and female control and iRAmKO mice were injected with tamoxifen for 5 consecutive days. After 2 weeks, the plantaris muscles were subjected to either a mechanical overload (MOV) or SHAM surgery. After 16 days, the plantaris muscles were collected, cross-sectioned at the mid-belly, and analyzed with immunohistochemistry for dystrophin. (**B**) Representative images from male mice of mid-belly cross-sections at two different levels of magnification. Scale bars = 200 µm for the low magnification images and 50 µm for the high magnification images. Measurements of the (**C**) plantaris muscle mass to body mass ratio, (**D**) whole muscle cross-sectional area (CSA), (**E**) number of fibers per cross-section (CS), and (**F**) average fiber CSA per muscle. All values are presented as group means ± SEM and expressed relative to the group mean of the sex-matched Control-SHAM samples, n = 10-13 / group. The data were analyzed with two-way ANOVA. * Significant effect of surgery within the given genotype. † Significant effect of genotype within the given surgical condition, P ≤ 0.05.

Following a 16-day recovery period, the PLTs were harvested and analyzed for changes in mass and whole muscle cross-sectional area (CSA). The outcomes revealed that MOV led to a robust increase in both the muscle-to-body mass ratio and mid-belly CSA, with the latter occurring through a partially mTORC1-dependent mechanism (Fig. 1B-D). As we have previously shown, the increase in mid-belly CSA can be attributed to the radial growth of the individual fibers as well as an increase in the number of fibers per cross-section (likely due to mTORC1-independent longitudinal growth of fibers, for details see ^7,13,22,24^). Therefore, we quantified the number of fibers per cross-section and found that the MOV-induced increase was mediated through an mTORC1-independent mechanism (Fig. 1E). We then measured the average CSA of the fibers, and in stark contrast, we found that the MOV-induced increase in fiber CSA was substantially reduced in the iRAmKO mice (Fig. 1F). Thus, in agreement with other recently published work ^22^, these data suggest that the modest increase in muscle-to-body mass ratio and whole muscle CSA in iRAmKO mice arises from an mTORC1-independent longitudinal growth of muscle fibers, whereas the induction of the radial growth of muscle fibers is mediated through a mechanism that is heavily dependent on mTORC1.

### MOV Induces Myofibrillogenesis and the Radial Growth of Myofibrils via an mTORC1-dependent mechanism

Having established that radial growth of muscle fibers is largely governed by an mTORC1-dependent mechanism, we then set out to test our hypothesis that mTORC1 is equally critical for the induction of myofibrillogenesis and the radial growth of myofibrils. Specifically, mid-belly cross sections of PLTs were subjected to immunostaining for dystrophin, to identify the periphery of muscle fibers, and SERCA1, which is enriched in the sarcoplasmic reticulum that surrounds individual myofibrils (Fig. 2A). For these analyses, a randomly selected subset of fibers from each sample were included, and as shown in Fig. 2B, the CSA of these fibers closely resembled the values observed for the entire population (see Fig. 1F and Fig. S1 along with methods for further details). Next, we applied our previously described FIM-ID pipeline to quantify the number of myofibrils per fiber and the average CSA of myofibrils ^18^. As shown in Fig. 2C-D, MOV significantly increased both the average number of myofibrils per fiber and the average myofibril CSA in muscles from the control mice. However, these MOV-induced adaptations were markedly reduced in the muscles from the iRAmKO mice. In other words, the results of our analyses indicate that mTORC1 significantly contributes to the induction of myofibrillogenesis and radial growth of myofibrils that occurs in response to MOV.

**Figure 2.**
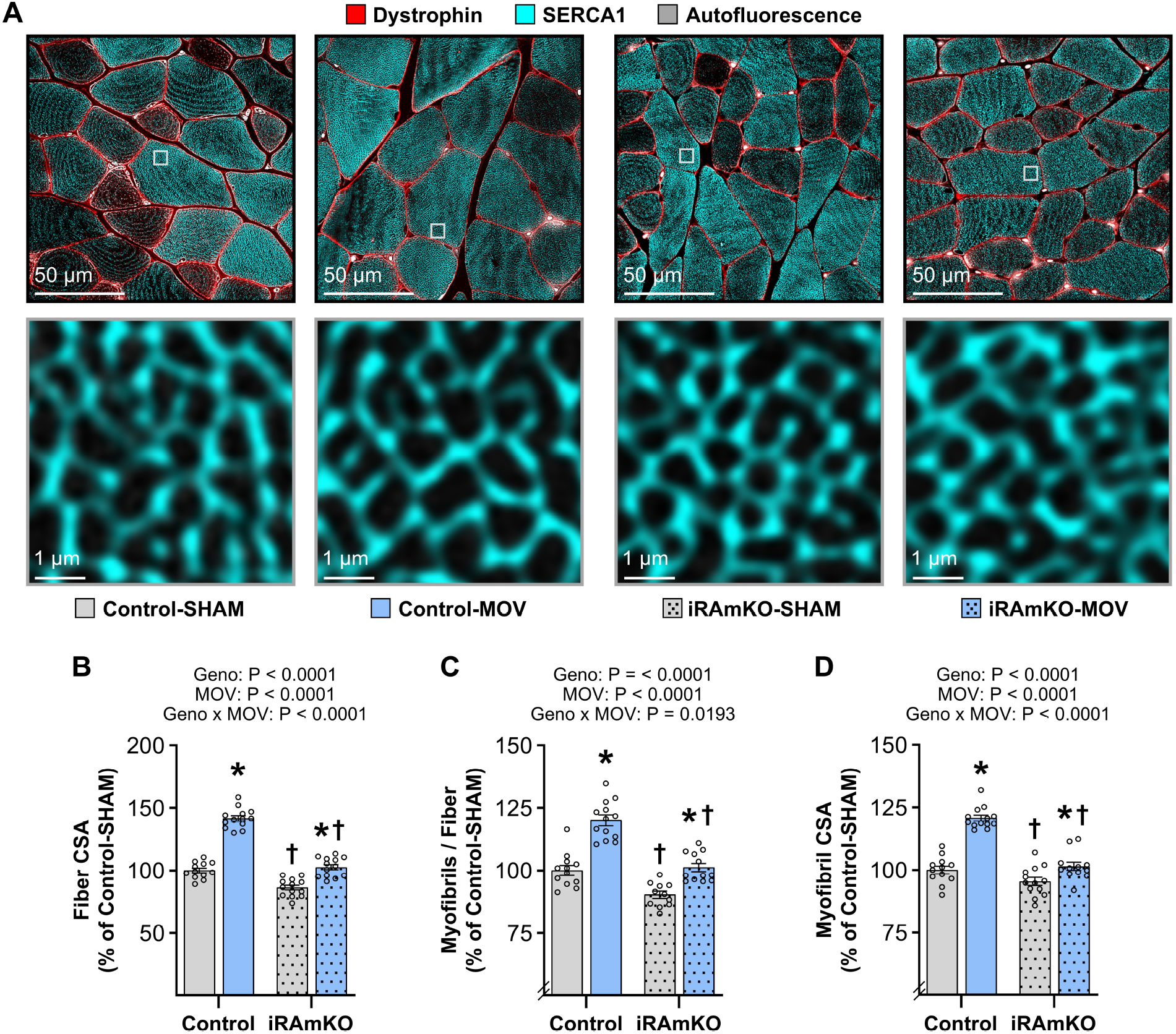
Mechanical Overload Induces Myofibrillogenesis and the Radial Growth of Myofibrils via an mTORC1-Dependent Mechanism. Male and female tamoxifen-treated control and iRAmKO mice were subjected to either a mechanical overload (MOV) or SHAM surgery. After 16 days, the plantaris muscles were collected, cross-sectioned at the mid-belly, and subjected to immunohistochemistry for dystrophin and SERCA1. The sections were then imaged with FIM-ID and merged with the signal from autofluorescence. (**A**) Representative images from male mice at two different levels of magnification. Scale bars = 50 µm for the low magnification images and 1 µm for the high magnification images. Measurement of the (**B**) cross-sectional area (CSA) of the fibers that were used to determine the (**C**) average myofibril CSA, and (**D**) the average number of myofibrils per fiber. All values are presented as group means ± SEM and expressed relative to the group mean of the sex-matched Control-SHAM samples, n = 10-13 / group. The data were analyzed with two-way ANOVA. Significant effect of surgery within the given genotype. † Significant effect of genotype within the given surgical condition, P ≤ 0.05.

### MOV Induces an Increase in Sarcomere Volume via an mTORC1-dependent mechanism

As noted above, recent studies have shown that MOV induces longitudinal growth of muscle fibers ^22^, a point that warrants careful consideration given the collection procedure employed in this study. Specifically, during the collections, the hindlimbs of all animals were fixed with the knee and ankle positioned at 90° angles. This is important because, if the entire muscle had undergone longitudinal growth, the imposed joint angles would likely force the fibers into an artificially shortened configuration. According to Poisson’s effect, such shortening would be expected to result in an increase in the CSA of the fibers as well as their respective myofibrils ^25^. To further appreciate this, myofibrils can be considered as a set of in-series connected sarcomeres, and the individual sarcomeres can be viewed as fluid-filled cylinders. According to the Poisson effect, when such cylinders are compressed/shortened, their cross-sectional area will expand. Thus, we wanted to determine whether the increase in myofibril CSA observed in Figure 2D might have simply been the result of an artifact of the collection procedure. To test this, once mid-belly cross-sections had been obtained, the samples were reoriented so that longitudinal sections could be derived from the same mid-belly region. The longitudinal sections were then subjected to immunostaining for alpha-actinin to demarcate the Z-discs of the sarcomeres and enable precise measurements of sarcomere length (Fig. 3A). In addition, as part of our standard collection procedure, whole-muscle lengths were recorded at the end of the fixation procedure, and as shown in Figure 3B, no significant differences in muscle length were observed between the experimental groups. Further, no significant differences in sarcomere length were observed across the experimental groups (Fig. 3A, C). We then used the average myofibril CSA from the analyzed samples to calculate sarcomere volume, and the outcomes revealed that both CSA and volume were significantly increased in the control-MOV group, and this effect was abolished by the loss of mTORC1 (Fig. 3D-E). Thus, it can be concluded that the observed differences in myofibril CSA were not an artifact of the collection procedure but rather were reflective of genuine mTORC1-dependent radial growth of the myofibrils.

**Figure 3.**
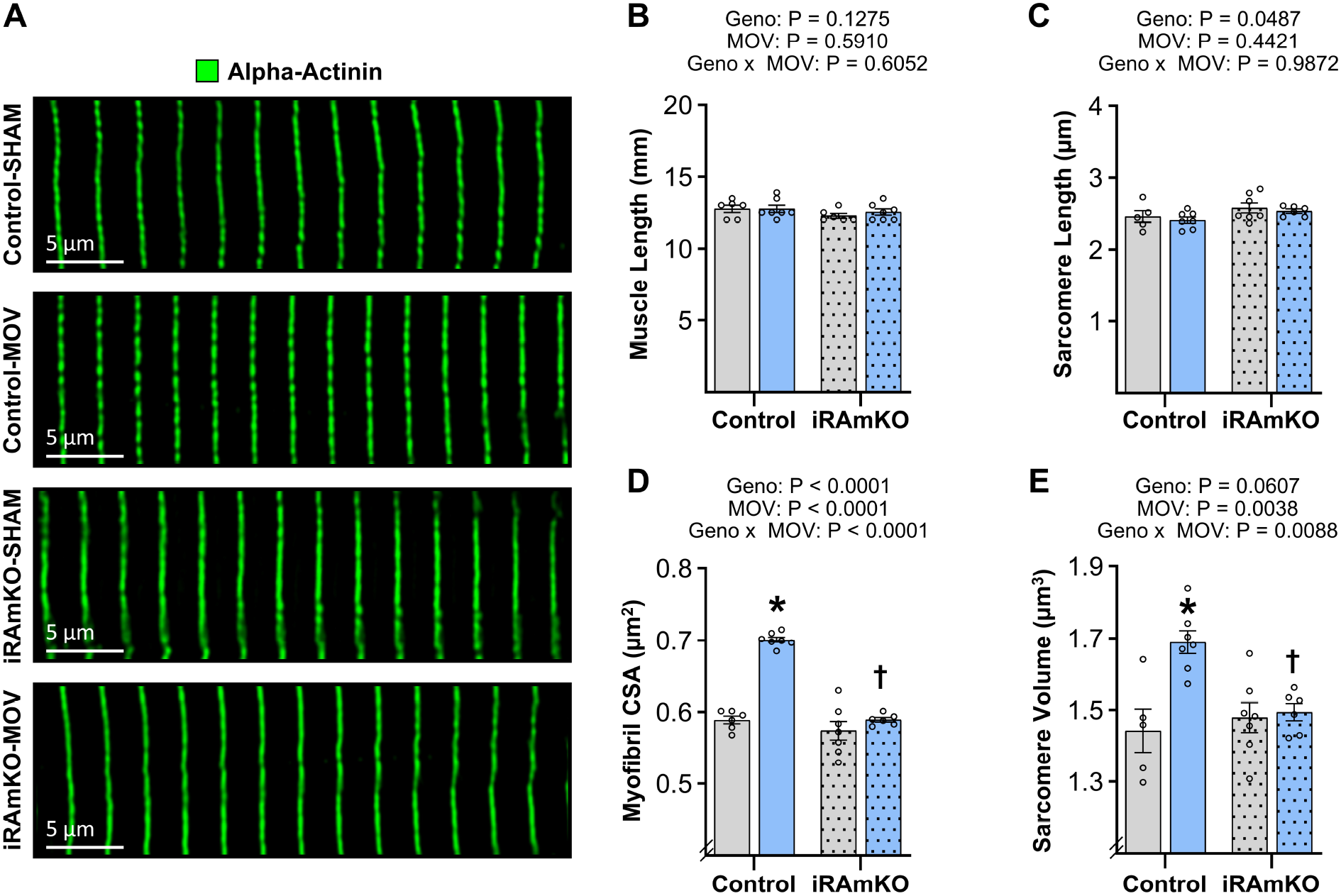
Mechanical Overload Induces an Increase in Sarcomere Volume via an mTORC1-Dependent Mechanism. Male tamoxifen-treated control and iRAmKO mice were subjected to either a mechanical overload (MOV) or SHAM surgery. After 16 days, the plantaris muscles were collected, cross-sectioned at the mid-belly, subjected to immunohistochemistry for dystrophin and SERCA1, and then imaged with FIM-ID to obtain measurements of myofibril cross-sectional area (CSA). (**A**) The same muscles were then longitudinally sectioned at the mid-belly and subjected to immunohistochemistry for alpha-actinin to obtain measurements of sarcomere length. Scale bars = 5 µm. Measurements of the (**B**) average muscle length, (**C**) average sarcomere length, (**D**) average myofibril CSA, and (**E**) average sarcomere volume. All values are presented as group means ± SEM, n = 5-7 / group. The data were analyzed by two-way ANOVA. * Significant effect of surgery within the given genotype. † Significant effect of genotype within the given surgical condition, P ≤ 0.05.

## Discussion

For decades, the structural adaptations of myofibrils and the mechanisms governing their response to increased mechanical loading have remained poorly defined. This is largely due to the technical challenges of electron microscopy and the labor-intensive manual tracing that is required to quantify myofibrils. In this study, we overcame these barriers through the use of our recently developed FIM-ID pipeline. By applying FIM-ID to both male and female mice, we also expanded upon previous studies that have relied exclusively on males. Specifically, our results did not show any significant interactions between the effects of sex, genotype, and MOV, suggesting that the growth response to increased mechanical loading is conserved across sexes. Moreover, our analyses enabled us to conclude that mTORC1 is a critical regulator of the myofibrillogenesis and radial growth of myofibrils that occur in response to mechanical overload.

A central question arising from these findings, is how mTORC1 controls these events in response to mechanical loading. Current dogma holds that growth is driven by a shift in the balance of protein synthesis and degradation towards a net increase in protein synthesis, allowing for the accumulation of newly synthesized proteins and subsequent growth. To this point, mTORC1 has long been appreciated for its role in the regulation of protein synthesis ^7,20,26-28^, and it has been widely assumed that the mechanical activation of mTORC1 drives muscle growth by stimulating protein synthesis ^21,27,29^. However, the activation of mTORC1 can also suppress protein degradation ^30,31^. Moreover, several recent studies have demonstrated that the loss of signaling through mTORC1 does not prevent the mechanically induced increase in protein synthesis ^7,22^. Collectively, this raises a key question: how does mTORC1 regulate radial growth if it’s not required for the increase in protein synthesis that is thought to drive the growth response?

To address this, we developed the model shown in Figure S2. Specifically, the model proposes that MOV induces an increase in protein synthesis via an undetermined mechanism, while concurrently activating mTORC1 ^22^, which functions as a suppressor of protein degradation and thereby allows for the net accumulation of proteins. These proteins can then be used in one of two processes: i) the formation of new myofibrils, or ii) incorporation into the pre-existing myofibrils, where they drive radial growth of the myofibrils. Importantly, as depicted in the model, when signaling through mTORC1 is inhibited, its suppressive effect on protein degradation is lost. As a result, the MOV-induced increase in protein synthesis would be offset by an elevation in protein degradation, and therefore, a net accumulation of proteins would not occur. Under this condition, new myofibrils could still be formed, but their formation would be countered by the accelerated breakdown of existing myofibrils, resulting in no overall increase in myofibril number. Similarly, newly synthesized proteins could still be incorporated into existing myofibrils, but radial growth of the myofibrils would not occur because the enhanced accumulation of newly synthesized proteins would be offset by the increased rate of protein degradation. Moving forward, it will be vital to test this hypothesis as it has the potential to redefine the role that mTORC1 plays in the mechanically induced growth of skeletal muscle.

Although our study strongly indicates that mTORC1 is crucial for myofibrillogenesis and the radial growth of myofibrils, there are some limitations that need to be acknowledged. For instance, due to the nature of the MOV procedure, our primary outcomes were restricted to the plantaris muscle. Since the plantaris is almost exclusively composed of fast-twitch fibers, it remains to be determined whether the same observations would be made in slow-twitch fibers. Furthermore, MOV induces a chronic mechanical stimulus and, therefore, the adaptations that it induces may not reflect those which occur in response to intermittent forms of mechanical stimuli such as resistance exercise. Thus, additional studies that extend the analyses to slow-twitch fibers and intermittent models of mechanical loading will be needed to fully define the extent to which mTORC1 governs the myofibril adaptations.

In summary, the outcomes of this study establish mTORC1 as a central regulator of the ultrastructural adaptations that drive the mechanically induced growth of skeletal muscle. Specifically, we reveal that the radial growth of muscle fibers is driven by myofibrillogenesis and the radial growth of myofibrils, and that both of these processes are mediated through an mTORC1-dependent mechanism. When combined with other recent developments in the field, we propose a model in which mTORC1 acts as a gatekeeper that permits the retention, rather than the synthesis, of proteins that drive the ultrastructural adaptations. Elucidating the molecular events through which mTORC1 exerts this control will be critical for developing targeted interventions to preserve or restore muscle mass in conditions such as sarcopenia, cancer cachexia, and prolonged bed rest.

## Methods

### Animals

iRAmKO mice have been previously described ^7^. At 6 weeks of age, male and female iRAmKO mice and littermate controls were injected intraperitoneally with a solution containing 2 mg tamoxifen every 24 h for 5 days as previously described ^7^. Mice were maintained at 25 °C on a 12-h light/dark cycle (lights on at 6:00 AM) with ad libitum access to standard chow and water. Two weeks after being treated with tamoxifen, the mice were randomly allocated to experimental groups and received surgical procedures under anesthesia. For tissue harvesting, animals were euthanized under anesthesia followed by cervical dislocation, and hindlimbs were subsequently collected. All experimental protocols were reviewed and approved by the University of Wisconsin–Madison Institutional Animal Care and Use Committee (IACUC; protocol #V005375).

### Mechanical Overload

Bilateral mechanical overload (MOV) procedures were carried out between 8:00 AM and 12:00 PM in accordance with our previously published methods ^18^. Briefly, mice were anesthetized using 1–5% isoflurane in oxygen, after which mechanical overload of the plantaris muscle (PLT) was induced by excising the distal one-third of the gastrocnemius muscle while preserving the soleus and plantaris muscles. Control animals underwent a bilateral sham operation consisting of a lower-leg incision that was subsequently closed without muscle removal. Surgical incisions were closed using non-absorbable nylon sutures (Oasis) in combination with Vetbond tissue adhesive (3M). Immediately after the procedure, mice were given an IP injection containing 0.05 μg/g of buprenorphine to alleviate any pain. Mice were also immediately placed on a rodent chow containing 500 mg/g tamoxifen (Envigo Teklad) and were maintained on this chow throughout the recovery period. Following 16 days of recovery, the mice were re-anesthetized with 1–5% isoflurane in oxygen, and the PLTs were harvested and processed as described below. Consistent with the surgical procedures, all terminal tissue collections were conducted between 8:00 AM and 12:00 PM.

### Collection and processing of mouse skeletal muscles

Hindlimb harvesting and tissue preparation were conducted as detailed previously ^18^. In brief, mice were anesthetized with 2–5% isoflurane in oxygen, after which the hindlimb skin was reflected and the limbs were disarticulated at the hip. For each limb, the foot and proximal femur were secured to a circular aluminum wire mesh with non-absorbable silk sutures, positioning the foot–tibia and tibia–femur joints at 90° angles. The mounted hindlimbs were then fixed in 20 mL of 4% paraformaldehyde in 0.1 M phosphate buffer (PB) for 3 h at room temperature on a tabletop rocker set to 50 RPM.

Following fixation, PLTs were dissected from each hindlimb. Muscle mass was measured, after which the PLTs were placed into 1.5 mL tubes containing 1.0 mL of 4% paraformaldehyde in 0.1 M PB and post-fixed for 21 h at 4 °C on a nutating rocker. Muscles were then cryoprotected by sequential incubation in 15% sucrose in 0.1 M PB for 6 h, followed by 45% sucrose in 0.1 M PB for 18 h at 4 °C. Finally, tissues were embedded in OCT compound (Tissue-Tek), rapidly frozen in isopentane cooled with liquid nitrogen, and stored at −80 °C until sectioning.

### Immunohistochemistry

Cross-sections (5 μm) from the mid-belly region of the PLTs were obtained with a cryostat maintained at −30 °C and mounted onto Superfrost Plus slides (Fisher Scientific). After cross-sections were obtained, the samples were reoriented to obtain longitudinal sections (3 μm) from the mid-belly region with the cryostat maintained at −35 °C, and again, the sections were mounted onto Superfrost Plus slides. In both instances, the mounted sections were immediately rehydrated in deionized water for 15 min. While keeping the sections hydrated, excess moisture surrounding each section was carefully removed with a Kimwipe, and a hydrophobic boundary was drawn around the section with a PAP pen (Aqua Hold 2, Scientific Device Laboratory #9804-02). Slides were then transferred to a humidified chamber, where all subsequent washes and incubations were conducted at room temperature on a rotating rocker set to 50 RPM.

Cross-sections and longitudinal sections were incubated for 30 min in a blocking solution consisting of 0.5% Triton X-100 and 0.5% bovine serum albumin in PBS. Cross-sections were then incubated overnight in blocking solution containing mouse IgG1 anti-SERCA1 (1:100, VE121G9, Santa Cruz #SC-58287) and rabbit anti-dystrophin (1:100, Thermo Fisher #PA1-21011), and longitudinal-sections were incubated overnight in blocking solution containing mouse IgG1 anti α-actinin (1:150, Novus Biologicals #NBP1-22630). Following primary antibody incubations, the sections were washed three times for 5-10 min in PBS, followed by three additional 0.5-1 hr washes in PBS. The cross-sections were then incubated overnight in blocking solution containing Alexa Fluor 594–conjugated goat anti-mouse IgG (Fcγ subclass 1–specific; 1:2000, Jackson ImmunoResearch #115-585-205) plus Alexa Fluor 488–conjugated goat anti-rabbit IgG (1:5000, Invitrogen #A11008), and longitudinal sections were incubated overnight in blocking solution containing Alexa Fluor 488–conjugated goat anti-mouse IgG1 (1:2000, Jackson ImmunoResearch 115-545-205). After secondary antibody incubation, the sections were washed three times for 5-10 min in PBS, followed by three additional 0.5-1 h washes in PBS. The washed cross-sections were mounted with 4 μL of ProLong™ Gold Antifade Mountant (Thermo Fisher #P36930), covered with a No. 1 glass coverslip and allowed to cure in the dark for a minimum of 24 h prior to imaging. The washed longitudinal sections were mounted with 4 μL of ProLong™ Diamond Antifade Mountant (Thermo Fisher #P36965), covered with a No. 1 glass coverslip and allowed to cure in the dark for a minimum of 48 h prior to imaging.

### Assessment of whole muscle size and muscle fiber size

Muscle cross-sections immunolabeled for dystrophin and SERCA1 were imaged using a Keyence BZX700 automated inverted epifluorescence microscope equipped with a 10× objective. For each fluorescence channel (FITC and TxRED), 3×3 tiled image fields were collected and stitched using Keyence Analyzer software. Whole-muscle cross-sectional area (CSA), as well as the average fiber CSA per muscle, were quantified using our previously established CellProfiler analysis pipeline ^32^.

### Fluorescence imaging of myofibrils with image deconvolution (FIM-ID)

FIM-ID was carried out as we have previously reported ^18^. Briefly, randomly selected regions of interest from dystrophin- and SERCA1-labeled cross-sections were imaged using a Leica THUNDER Imager Tissue 3D microscope equipped with an HC PL APO 63×/1.40 oil immersion objective. Z-stack images spanning the full thickness of the sections were acquired and processed using Leica’s Small Volume Computational Clearing (SVCC) deconvolution algorithm. The deconvoluted image stacks were subsequently imported into ImageJ, where the Z-plane exhibiting optimal SERCA1 focus was identified. The signal for autofluorescence from the corresponding plane was then merged with the SERCA1 signal using ImageJ’s Z-Project function, and the merged image was saved as a single grayscale TIFF file. Final merged images were converted to 16-bit TIFF format and resized to a standardized resolution of 6144 × 6144 pixels using ImageJ.

### Automated measurements of myofibril size and number with FIM-ID

Individual muscle fibers were randomly chosen from each FIM-ID image by an investigator blinded to experimental grouping and manually outlined in ImageJ to determine the fiber CSA, as detailed previously ^18^. Initial analyses included 15 autofluorescent and 15 non-autofluorescent fibers with additional fibers randomly selected until the mean CSA of the sampled fibers fell within ±7.5% of the average fiber CSA as derived from the Keyence-based whole-section analyses. Images of individual SERCA1-positive fibers were then analyzed using an automated CellProfiler pipeline (“Myofibril CSA Analysis”) in CellProfiler version 4.2.1 to determine the average CSA of the myofibrils ^18^. To determine the number of myofibrils per fiber, each SERCA1 fiber image was further processed using the “Intermyofibrillar Area” CellProfiler pipeline, which measures the area of the fiber that is occupied by intermyofibrillar components (e.g., sarcoplasmic reticulum and mitochondria). The intermyofibrillar area was subtracted from the total fiber CSA to calculate the myofibrillar area, which was subsequently divided by the average CSA of the myofibrils in that fiber.

### Determination of sarcomere length and volume

For each sample, 5 randomly selected regions of interest from α-actinin labeled longitudinal sections were imaged using a Leica THUNDER Imager Tissue 3D microscope equipped with an HC PL APO 63×/1.40 oil immersion objective. Z-stack images spanning the full thickness of the sections were acquired and processed using Leica’s Small Volume Computational Clearing (SVCC) deconvolution algorithm (refractive index set to 1.47, strength set to 80-100%, and regularization set to 0.05). The deconvoluted image stacks were subsequently imported into ImageJ, where the Z plane with the most in-focus image for α-actinin was selected. Next, within each image, four randomly selected regions had a line drawn perpendicular to the Z-discs. The lines were bound by the Z-discs (marked by peaks in α-actinin signal) and spanned 10-20 Z-discs in length. The number of Z-discs along the length of each line was determined and divided by the number of Z-discs minus 1 to obtain an average sarcomere length. A total of 20 measurements were obtained for each sample, and the mean of these measurements was recorded as the average sarcomere length. The average sarcomere volume was then calculated by multiplying the average sarcomere length for the sample by the same sample’s average myofibril CSA.

### Statistical analysis

Final data sets were compiled, and then samples that deviated more than three times from the mean within a given group were identified and excluded as outliers, as previously described ^18^. Statistical significance was then assessed using two-way ANOVA with Fisher’s LSD post hoc comparisons, or three-way ANOVA with Sidak’s post hoc comparisons as detailed in the figure legends. Differences were considered significant at P < 0.05. All analyses were conducted using GraphPad Prism version 10.6.1 (Windows).

## Figure Legends

**Supplemental Figure 1. The Role of mTORC1 in the Mechanical Overload-Induced Growth of Skeletal Muscle in Male and Female Mice**. Male and female tamoxifen-treated control and iRAmKO mice were subjected to either a mechanical overload (MOV) or SHAM surgery. After 16 days, the plantaris muscles were collected and subjected to measurements of the (**A**) muscle mass to body mass ratio, (**B**) whole muscle cross-sectional area (CSA), (**C**) number of fibers per cross-section (CS), (**D**) average fiber CSA, (**E**) average myofibril CSA, and (**F**) average number of myofibrils per fiber. All values are presented as group means ± SEM, n = 5-7 / group. The data were analyzed by three-way ANOVA. * Significant effect of sex within the given genotype and surgical condition, P ≤ 0.05.

**Supplementary Figure 2. Potential Mechanism for the Role of mTORC1 in Mechanically Induced Myofibrillogenesis and the Radial Growth of Myofibrils**. (**A**) The model predicts that under normal conditions, mechanical overload (MOV) induces an increase in protein synthesis through an as-yet undefined mechanism. Simultaneously, MOV activates mTORC1, which acts to suppress protein degradation and thereby permits the net accumulation of proteins. These proteins can then contribute to one of the processes that drive the radial growth of the muscle fibers: i) myofibrillogenesis via the formation of new myofibrils, or ii) radial growth of the myofibrils. (**B**) When signaling through mTORC1 is inhibited, the suppressive effect it exerts on protein degradation is lost, and the MOV-induced increase in protein synthesis is offset by the increased rate of protein degradation. Consequently, while new myofibrils are still formed, their formation is countered by the accelerated breakdown of existing myofibrils, resulting in no overall increase in myofibril number. Likewise, while newly synthesized proteins can still be incorporated into existing myofibrils, radial growth does not occur because the accumulation of these proteins is offset by the heightened rate of degradation.

## Acknowledgements

This work was supported by the National Institute of Arthritis and Musculoskeletal and Skin Diseases of the National Institutes of Health (NIH)under awards R01AR082816 to T.A.H and F31AR0085939 to C.G.K.F. Elements of supplementary figure 2 were created in BioRender. Hornberger, T. (2026) https://BioRender.com/rkrepdo

## Author Contributions

Conceptualization: CGKF, RKAS, TAH

Methodology: RKAS, WGZ, TAH

Investigation: RKAS, ANL

Formal Analysis: CGKF, RKAS

Visualization: CGKF, RKAS, TAH

Supervision: TAH

Writing – original draft: CGKF, TAH

Writing – reviewing & editing: CGKF, RKAS, ANL, WGK, TAH

## Resource Availability

All data are available upon request to the corresponding author.

