## Supplemental Figures for "Mechanical Loading Induces the Radial Growth of Myofibrils and Myofibrillogenesis via an mTORC1-Dependent Mechanism"

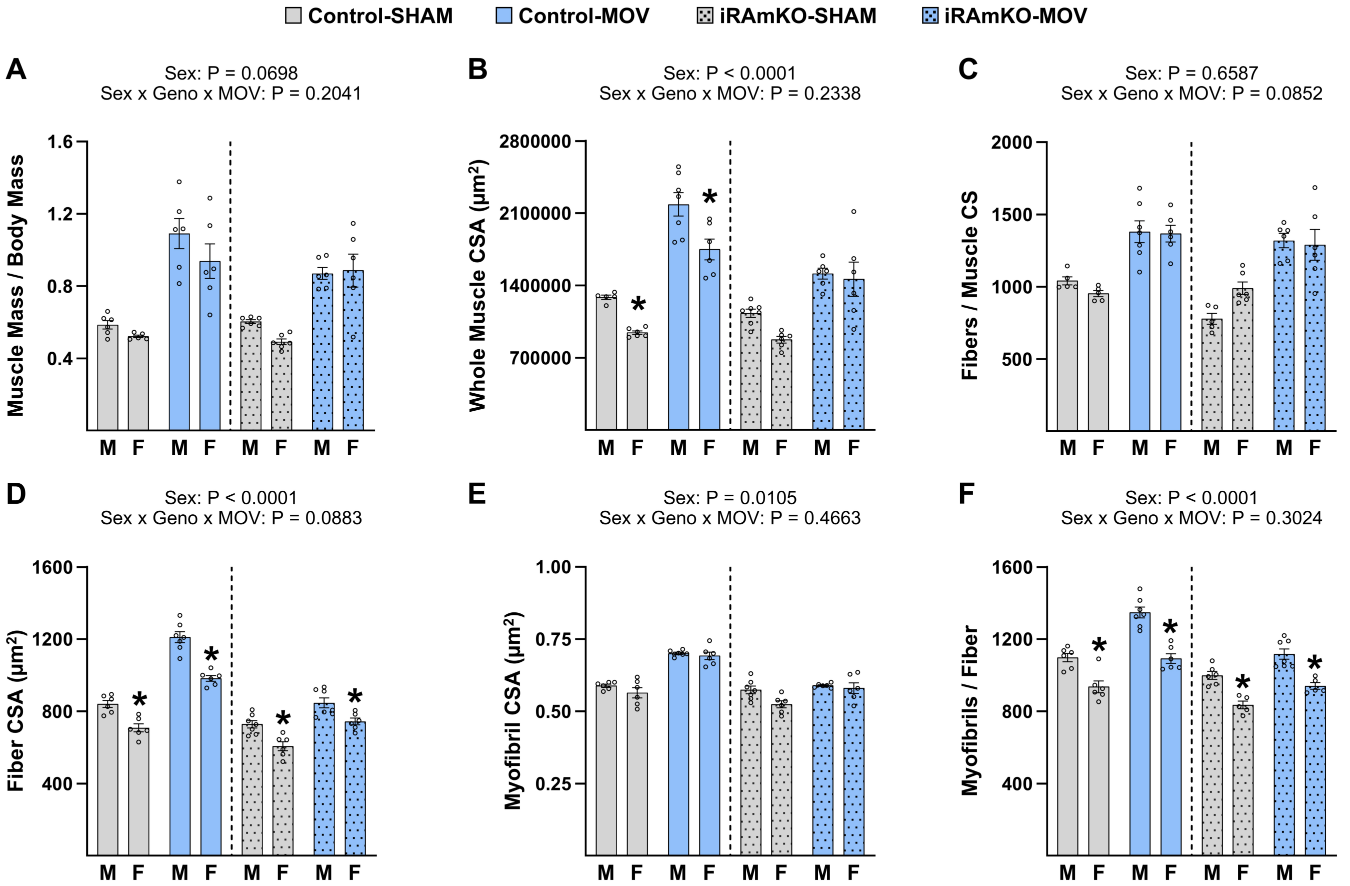


**Supplemental Figure 1. The Role of mTORC1 in the Mechanical Overload-Induced Growth of Skeletal Muscle in Male and Female Mice**. Male and female tamoxifen-treated control and iRAmKO mice were subjected to either a mechanical overload (MOV) or SHAM surgery. After 16 days, the plantaris muscles were collected, and subjected to measurements of the (**A**) muscle mass to body mass ratio, (**B**) whole muscle cross-sectional area (CSA), (**C**) number of fibers per cross-section (CS), (**D**) average fiber CSA, (**E**) average myofibril CSA, and (**F**) average number of myofibrils per fiber. All values are presented as group means ± SEM, n = 5-7 / group. The data were analyzed by three-way ANOVA. * Significant effect of sex within the given genotype and surgical condition, $P \leq 0.05$.

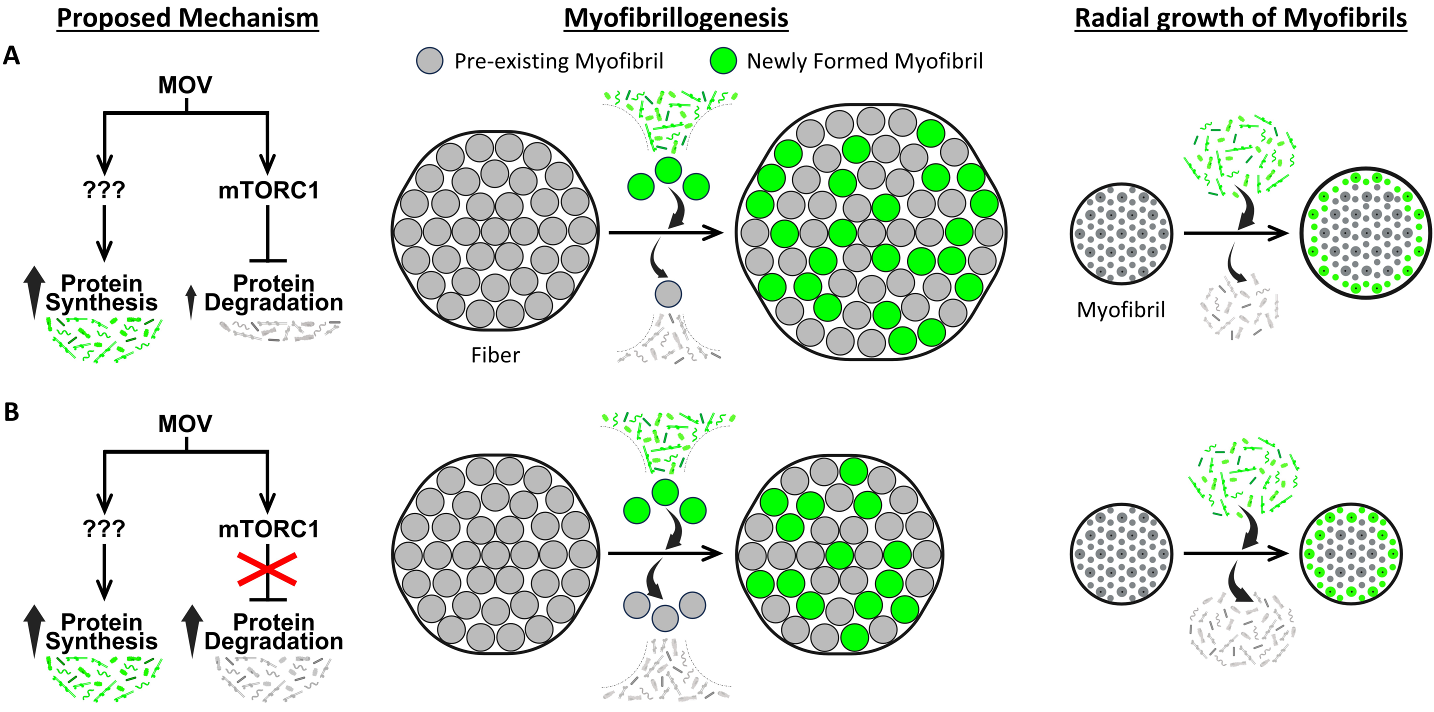


**Supplementary Figure 2. Potential Mechanism for the Role of mTORC1 in Mechanically Induced Myofibrillogenesis and the Radial Growth of Myofibrils.** (**A**) The model predicts that under normal conditions, mechanical overload (MOV) induces an increase in protein synthesis through an as-yet undefined mechanism. Simultaneously, MOV activates mTORC1, which acts to suppress protein degradation and thereby permits the net accumulation of proteins. These proteins can then contribute to one of the processes that drive the radial growth of the muscle fibers: i) myofibrillogenesis via the formation of new myofibrils, or ii) radial growth of the myofibrils. (**B**) When signaling through mTORC1 is inhibited, the suppressive effect it exerts on protein degradation is lost, and the MOV-induced increase in protein synthesis is offset by the increased rate of protein degradation. Consequently, while new myofibrils are still formed, their formation is countered by the accelerated breakdown of existing myofibrils, resulting in no overall increase in myofibril number. Likewise, while newly synthesized proteins can still be incorporated into existing myofibrils, radial growth does not occur because the accumulation of these proteins is offset by the heightened rate of degradation.
